# Ventral Tegmental Area Glutamatergic Neurons Suppress Hippocampal Sharp-wave Ripples and Induce Head Movements in Mice

**DOI:** 10.1101/2025.11.23.690012

**Authors:** Manivannan Subramaniyan, Sumithrra Manivannan, Maggie S. Zhou, John A. Dani

## Abstract

Sharp-wave ripples are brief, high-frequency hippocampal oscillations crucial for memory consolidation, occurring predominantly during the non-rapid-eye-movement (NREM) sleep. While ripples are generated intrinsically within the hippocampus, growing evidence suggests that their occurrence is modulated by subcortical regions. Whether the ventral tegmental area (VTA)—a subcortical region that directly projects to the hippocampus—regulates ripple activity remains unknown. To investigate this, we optogenetically activated VTA neurons during NREM sleep and recorded ripple activity in the dorsal hippocampus via field recording electrodes. Using cell-type specific excitatory opsin expression in the VTA, we found that activation of glutamatergic (vGlut2+) neurons strongly suppressed ripple incidence. In contrast, activation of GABAergic (vGAT+) neurons produced weak suppression, and activation of dopaminergic (DAT+) neurons showed no effect. During ripple suppression by glutamatergic activation, we observed small but consistent head movements. Trial-by-trial analysis revealed no correlation between head movement and ripple suppression. Furthermore, glutamatergic activation during wakefulness also led to head movement, suggesting that ripple suppression and head movement could be dissociated. Overall, our results suggest that VTA vGlut2+ neurons play a role in head movement and that their activation suppresses hippocampal sharp-wave ripples during NREM sleep, offering a potential way to alter memory consolidation.

## Introduction

The hippocampus is essential for learning, memory, and spatial navigation, and it exhibits distinct electrophysiological patterns across behavioral states. During behavioral immobility and non-rapid-eye-movement (NREM) sleep, brief (∼100 ms), high-frequency (∼200 Hz) oscillations super-imposed on large deflections in the local field potential are readily observed in the hippocampus [1, 2]. These oscillatory events, called sharp-wave rip-plesarise from intrinsic hippocampal activity [3-5] and provide a mechanism for transferring memories from the hippocampus to the neocortex [6-8]. Disruption of ripples during post-learning slow-wave sleep impairs spatial learning and memory [9-12], whereas prolonging their duration enhances memory [13]. Understanding how ripple occurrence is regulated is therefore critical, offering potential therapeutic avenues to restore memory function or attenuate maladaptive memories, such as those in post-traumatic stress disorder (PTSD).

Evidence from several studies indicates that cortical and subcortical brain regions connected to the hippocampus can influence the occurrence of hippocampal ripples. In mice, optogenetic silencing of the medial entorhinal cortex reduced ripple incidence in the hippocampus [14]. In non-human primates, combined fMRI and electrophysiology revealed that most subcortical regions—including the ventral tegmental area (VTA), locus coeruleus (LC) and raphe—were suppressed during hippocampal ripples [15], suggesting that increasing neural activity in these regions could suppress ripples. Supporting this idea, ripple suppression in the hippocampal CA1 region has been observed following optogenetic activation of median raphe neurons during NREM sleep in mice [16], electrical stimulation of the LC in rats [17], and optogenetic stimulation of medial septal cholinergic neurons in mice [18]. Whether the VTA can similarly modulate ripple occurrence remains unknown.

The VTA neuronal population consists primarily of dopaminergic (∼60%), GABAergic (∼35%) and glutamatergic neurons (∼5%) [19, 20], all of which directly project to the hippocampus [21-29]. To examine the role of the VTA in controlling hippocampal ripple incidence, we optogenetically activated different VTA cell types in mice during NREM sleep while simultaneously monitoring ripple incidence in the dorsal CA1 region. Because the VTA is extensively interconnected with brain regions involved in motivated behaviors [30], and recent studies have highlighted its role in movement [31-33], we also monitored animal behavior and assessed the relationship between VTA neuronal activation, ripple incidence, and movement.

## Results

To examine the influence of the VTA on hippocampal sharp-wave ripple occurrence, we activated VTA neurons using Channelrhodopsin-2 (ChR2) while monitoring ripples in the dorsal CA1 region using 4–6 implanted electrodes (Fig. 1A). Experiments were conducted in the home cage of the mice during the light phase of the daily cycle when the animals normally sleep (Fig. 1B). Ripple analysis was restricted to NREM sleep, during which ripples are far more frequent than rapid-eye-movement (REM) sleep. While continuously recording both animal movement and CA1 neural activity, we activated ChR2 in the VTA with brief blue light pulses delivered at random inter-pulse intervals of 15–20 s (Fig. 1C). Ripple events were detected offline in each recording electrode by thresholding the spectral power in the ripple frequency band (Fig 1D). A median of 2 electrodes per mouse (Interquartile Range, IQR: 2–3) contained detectable ripples, and a median of 145 photostimulation trials (IQR: 111–237) were collected per experimental condition per mouse. Opsin expression in the VTA region was confirmed by anti-GFP and anti-TH immunostaining, and electrode and optic fiber placements were verified histologically.

**Figure 1.**
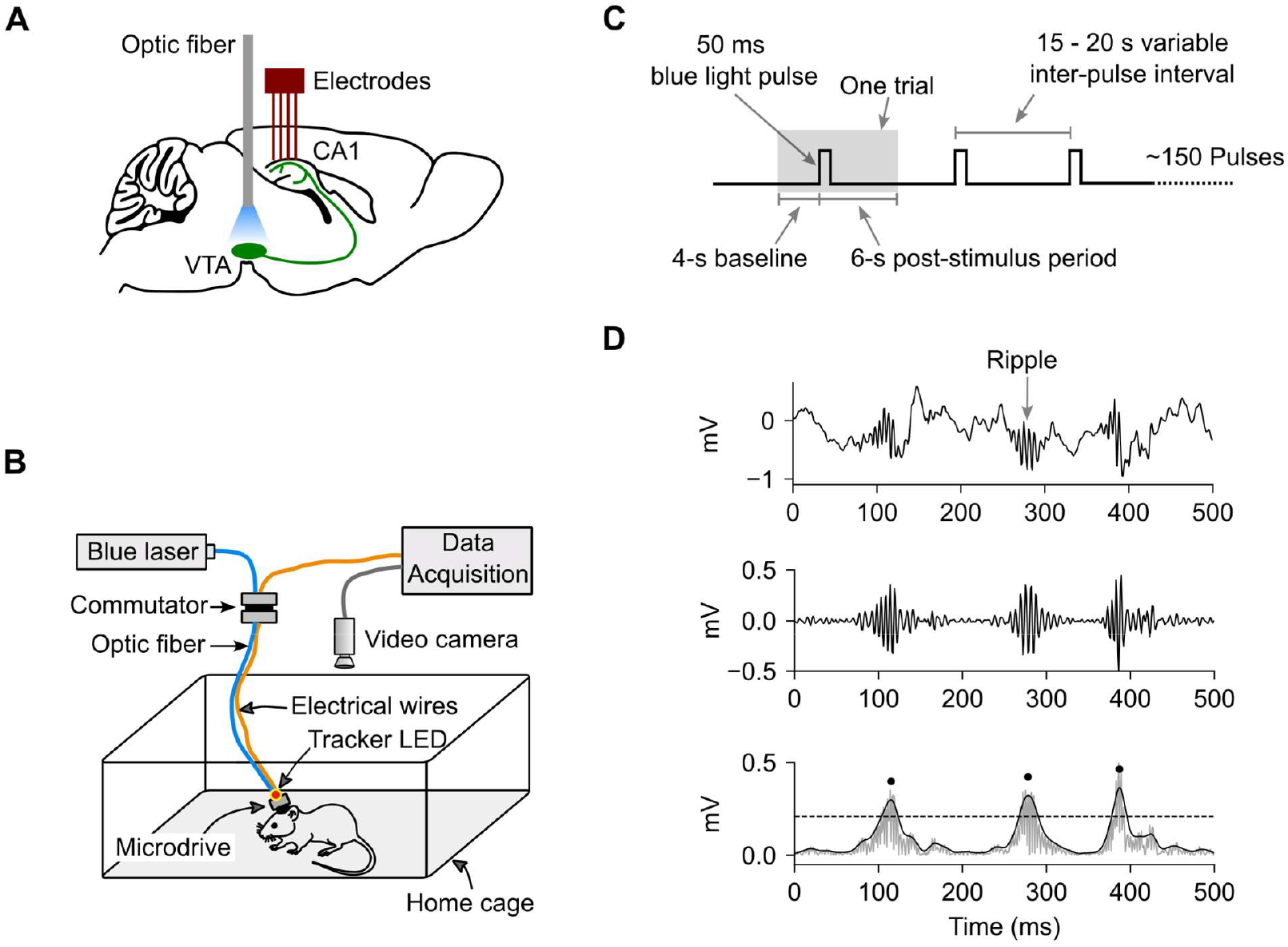
Experiment setup. **A**) Following viral injections to express opsins in the ventral tegmental area (VTA) neurons, an optic fiber was implanted in the VTA of either the left or right hemisphere, and a lightweight microdrive carrying 4–6 recording electrodes was implanted in the dorsal hippocampus ipsilateral to the optic fiber. **B**) Experiments were conducted in the mouse’s home cage (cage contents not shown for clarity) during the light (i.e., sleep) phase of the daily cycle. Cables for photostimulation and neural recording were connected to the head implant via a motorized commutator to prevent cable tangling. An overhead camera recorded animal movement. **C**) Photostimulation consisted of 50-ms light pulses delivered ∼150 times with a variable inter-stimulation interval of 15–20 s. A stimulation “trial” (shaded region) consisted of a 4-s pre-stimulus baseline and a 6-s post-stimulus period. **D**) To detect ripples, the raw data (top panel) was band-pass filtered (middle panel) and squared. The peaks of the envelope (bottom panel, dark trace) of the squared signals (gray trace) that crossed a pre-defined threshold (horizontal dashed line) were taken as ripple events (black filled circles).

We first expressed ChR2 in VTA neurons of wild-type mice without cell-type specificity. Figures 2A-2E show results from a representative mouse. Figure 2A shows ChR2 expression and the placement of the optic fiber and electrode tips. To examine the effect of VTA activation on ripple events, we plotted ripple times from each trial relative to light pulse onset (Fig. 2B). Brief activation of VTA neurons (5 ms) markedly reduced ripple occurrence in dorsal CA1 region for ∼1 s, well beyond the stimulus duration (Fig. 2B, C).

**Figure 2.**
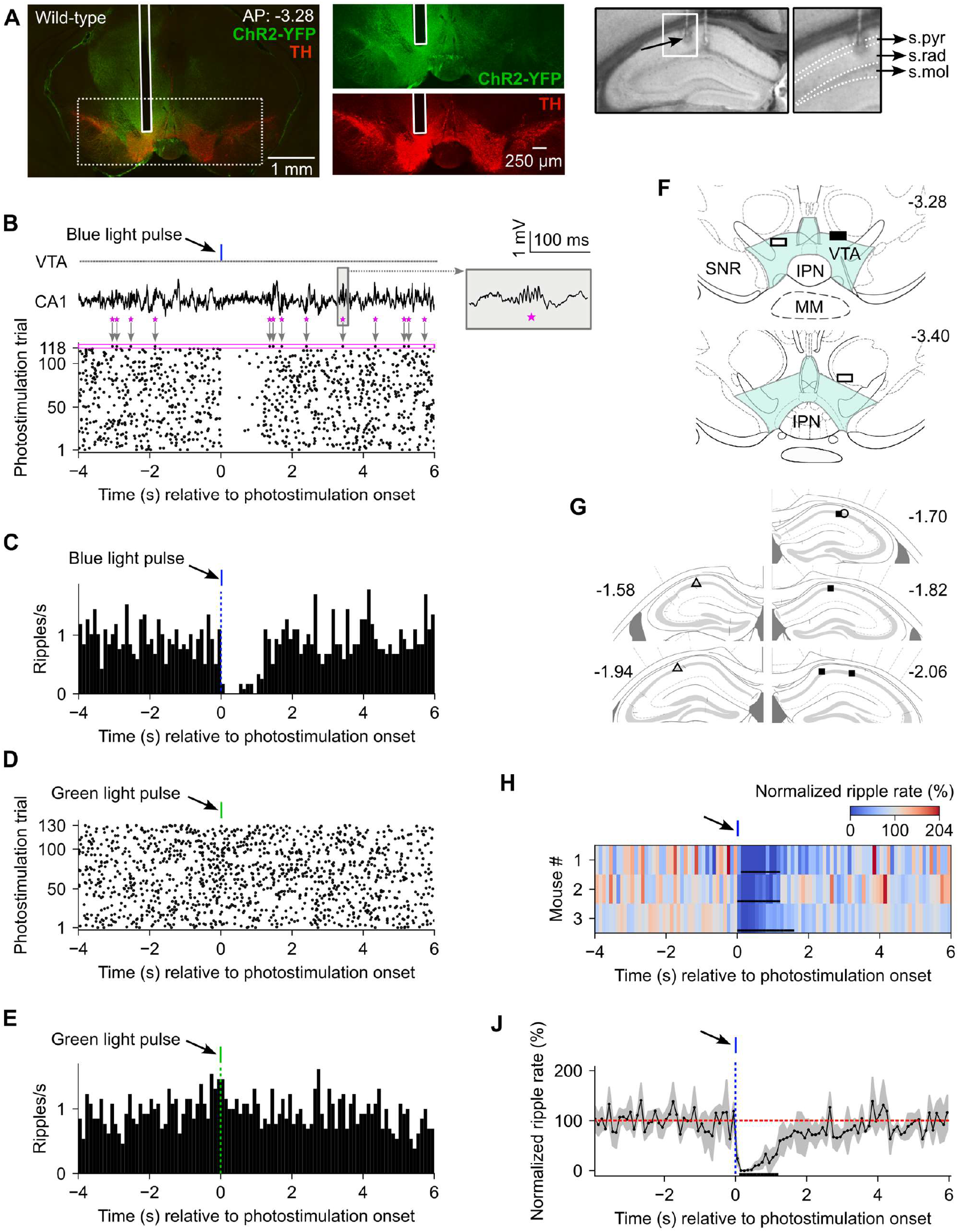
Effect of VTA stimulation on ripple incidence in wild-type mice during NREM sleep. **A**) Left: Immunostaining showing overlay of the expression of ChR2-YFP (green) and tyrosine hydroxylase (TH, red) in the VTA of an example mouse. The narrow vertical white rectangle marks the optic fiber track. The green and red channels within the dotted rectangle are expanded in the two adjacent panels. Right: Grayscale images showing two of the electrodes and lesion marks from the same mouse. Data from the electrode marked by the black arrow are shown in panels B–E. The region outlined by the white rectangle is expanded on the right, with the hippocampal layers indicated [stratum pyramidale (s.pyr), stratum radiatum (s.rad), and stratum moleculare (s.mol)]. **B**) Bottom: Raster plot of ripple events (black dots) aligned to light-pulse onset (time zero). A single trial is highlighted by the magenta box, enclosing several ripple events. Top: Photostimulation timeline (gray line labeled “VTA”), with the blue tick indicating a 50-ms light pulse. Raw data from the magenta-boxed trial are shown below, with ripples marked by magenta stars. One ripple (gray rectangle) is expanded in the inset on the right. **C**) Peristimulus time histogram of ripple rate. **D & E**) Raster plot (D) and peri-stimulus time histogram (E) of ripple rate for VTA stimulation with 50-ms green light (i.e., control) pulses. **F**) Optic fiber tip locations (rectangles), with fill color indicating sex (black = male, white = female). Blue shaded region indicates the VTA, with nearby structures labeled (IPN, interpeduncular nucleus; MM, medial mammillary nuclei; SNR, substantia nigra pars reticulata). **G**) Hippocampal electrode tip locations, with marker type distinguishing mice and fill color indicating sex (black = male, white = female). **H**) Pseudocolor plot of normalized ripple rate (%). Red indicates ripple rates above baseline; blue indicates ripple rates below baseline (suppression). Black horizontal bars mark periods of significant modulation. **J**) Normalized ripple rate averaged across mice (n = 3). Black horizontal bar marks the time window of significant modulation common for all mice. Shaded area indicates ± 1 standard deviation.

To test whether this suppression was due to heat from the light pulses rather than ChR2 activation, we delivered green light pulses, which generate equal or greater heat but minimally activate ChR2. Green light delivery in the same mouse did not sup-press ripples (Fig. 2D, E), indicating that the suppression observed with blue light (Fig. 2B, C) was due to ChR2-mediated neuronal activation.

We repeated the blue-light stimulation experiment in two additional mice, with the placement of the optic fiber and electrode tips shown in Figures 2F and 2G respectively. For population analysis, we normalized each mouse’s trial-averaged post-stimulus ripple rate to its baseline rate (Fig. 2H) and then averaged across mice (Fig. 2J). Because of the small sample size (n = 3), we did not conduct population-level statistical testing. However, within each mouse a cluster-size permutation test (see Methods) revealed significant ripple suppression lasting ∼1.2 s after stimulus onset (n = 51–244 trials per mouse; *p* < 0.05), demonstrating that VTA activation strongly suppresses dorsal hippocampal ripples.

The VTA predominantly contains dopaminergic (DA), GABAergic, and glutamatergic neurons, all of which project to the hippocampus. To assess cell-type contributions to ripple suppression, we expressed ChR2 selectively in DA neurons of DAT-Cre mice (Fig. 3A and Fig. S1). In an example mouse, activation of ChR2 in DA neurons with a single 50-ms blue-light pulse did not measurably alter ripple occurrence (Fig. 3B, C). At the population level, ripple rate remained unchanged (n = 6 mice; *p* > 0.05, cluster-size permutation test; Fig. 3D, E). Increasing VTA stimulation to 5-s stimulus trains in a subset of these mice likewise failed to produce a significant change in ripple rate (n = 5 mice; *p* > 0.05, cluster-size permutation test; Fig. S2E–H). Together, these results suggest that activation of VTA dopaminergic neurons does not significantly or measurably suppress hippocampal ripples.

**Figure 3.**
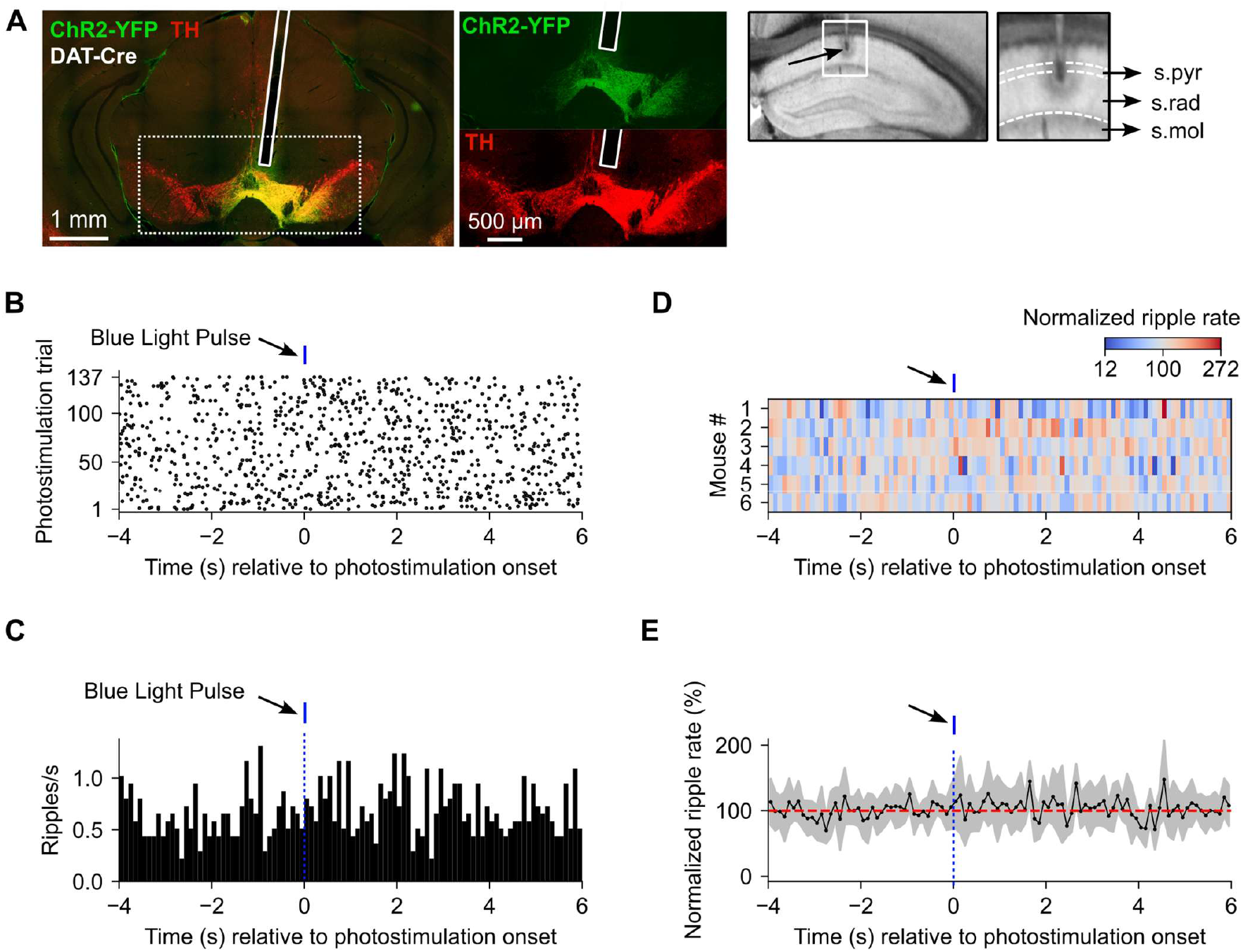
Effect of VTA stimulation on ripple incidence in DAT-Cre mice during NREM sleep. **A**) Left: Immunostaining showing overlay of the expression of ChR2-YFP (green) and tyrosine hydroxylase (TH, red) in the VTA of an example mouse. The narrow vertical white rectangle marks the optic fiber track. The green and red channels within the dotted rectangle are expanded in the two adjacent panels. Right: Grayscale images showing lesion marks of an electrode from the same mouse, with the electrode tip indicated by the black arrow. Data from this electrode are shown in panels B and C. **B**) Raster plot of ripple events (black dots) aligned to the onset of a 50-ms stimulation light pulse (blue tick). **C**) Peri-stimulus time histogram of ripple rate. **D**) Pseudocolor plot of normalized ripple rate (%) for the mouse population (n = 6). Red indicates ripple rates above baseline; blue indicates ripple rates below baseline (suppression). **E**) Normalized ripple rate averaged across mice (n = 6). Shaded area indicates ± 1 standard deviation. No significant modulation was detected either at the individual mouse level (panel D) or at the population level (panel E).

Next, because VTA activation in wild-type mice decreased rather than increased ripple occurrence, we hypothesized that VTA inhibitory neurons might mediate this effect. Therefore, we expressed ChR2 in GABAergic neurons of vGAT-Cre mice (Fig. 4A, Fig. S3). In an example mouse, activating VTA GABAergic neurons using a single 50-ms blue light pulse produced only mild ripple suppression (Fig. 4B, C). At the population level (Fig. 4D, E), a slight dip in the ripple rate was not significant (n = 6 mice; *p* > 0.05, cluster-size permutation test; Fig. 4E). To test whether stronger stimulation would enhance the effect, we delivered 5-s trains of blue-light pulses in a subset of these mice, but again we observed no significant modulation of ripple rate (n = 5; *p* > 0.05, cluster-size permutation test; Fig. S4E, F).

**Figure 4.**
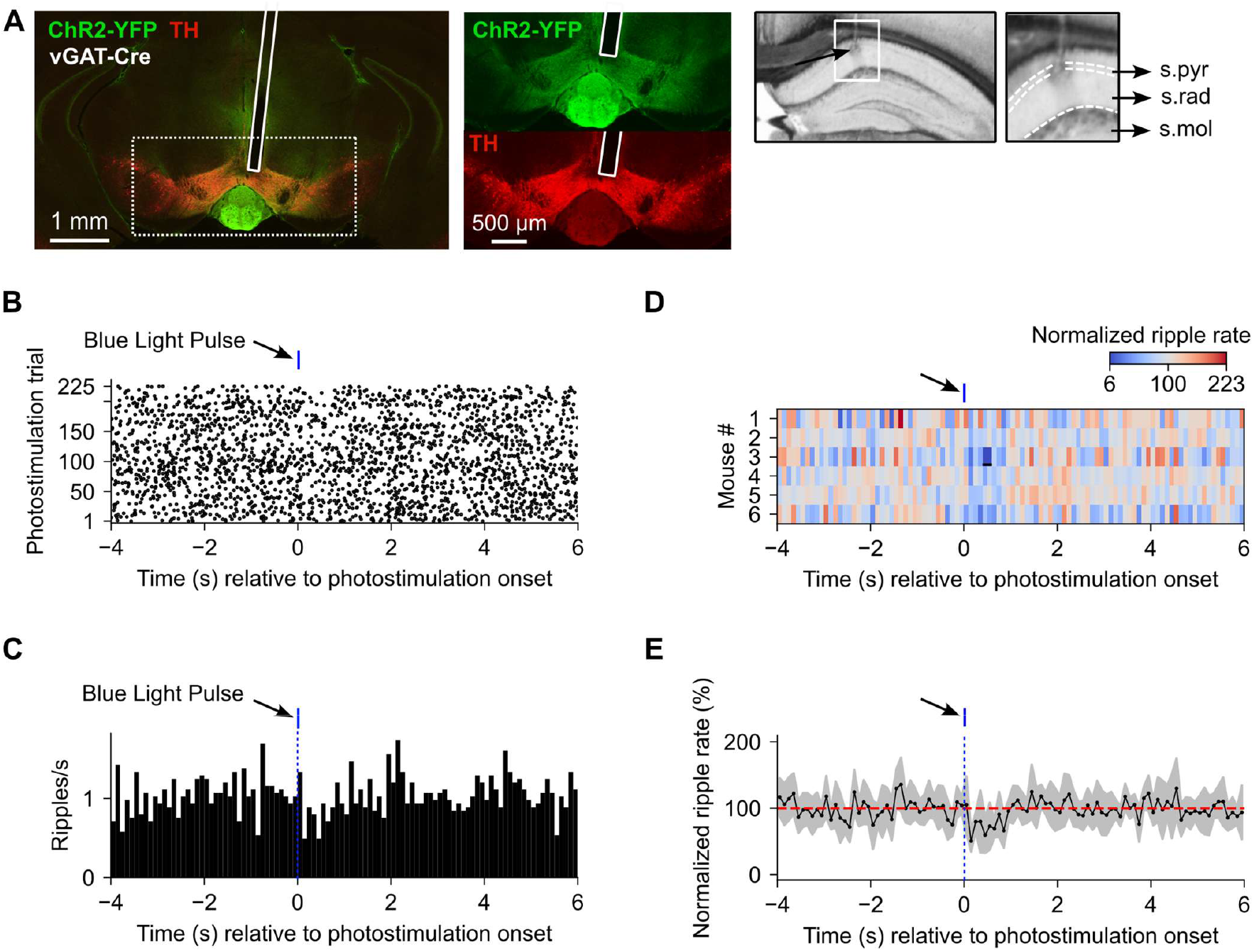
Effect of VTA stimulation on ripple incidence in vGAT-Cre mice during NREM sleep. **A**) Left: Immunostaining showing overlay of the expression of ChR2-YFP (green) and tyrosine hydroxylase (TH, red) in the VTA of an example mouse. The narrow vertical white rectangle marks the optic fiber track. The green and red channels within the dotted rectangle are expanded in the two adjacent panels. Right: Grayscale images showing lesion marks of an electrode from the same mouse, with the electrode tip indicated by the black arrow. Data from this electrode are shown in panels B and C. **B**) Raster plot of ripple events (black dots) aligned to the onset of a 50-ms stimulation light pulse (blue tick). **C**) Peri-stimulus time histogram of ripple rate. **D**) Pseudocolor plot of normalized ripple rate (%) for the mouse population (n = 6). The blue tick above the plot marks the 50-ms blue light pulse. Red indicates ripple rates above baseline; blue indicates ripple rates below baseline (suppression). Black horizontal bar marks a period of significant modulation in mouse 3. **E**) Normalized ripple rate averaged across mice (n = 6). Shaded area indicates ± 1 standard deviation. No significant modulation was detected at the population level.

The minimal ripple suppression observed with activation of GABAergic or DA neurons pointed to glutamatergic neurons as the most likely mediator of the strong suppression seen in wild-type mice. Indeed, in an example mouse, brief activation of ChR2-expressing vGlut2+ glutamatergic neurons in the VTA (Fig. 5A) produced an almost complete absence of ripples for ∼1 s (Fig. 5B, C), closely matching the effect in the wild-type mice. These findings indicate that VTA vGlut2+ neurons can powerfully suppress sharp-wave ripples in the dorsal hippocampus.

**Figure 5.**
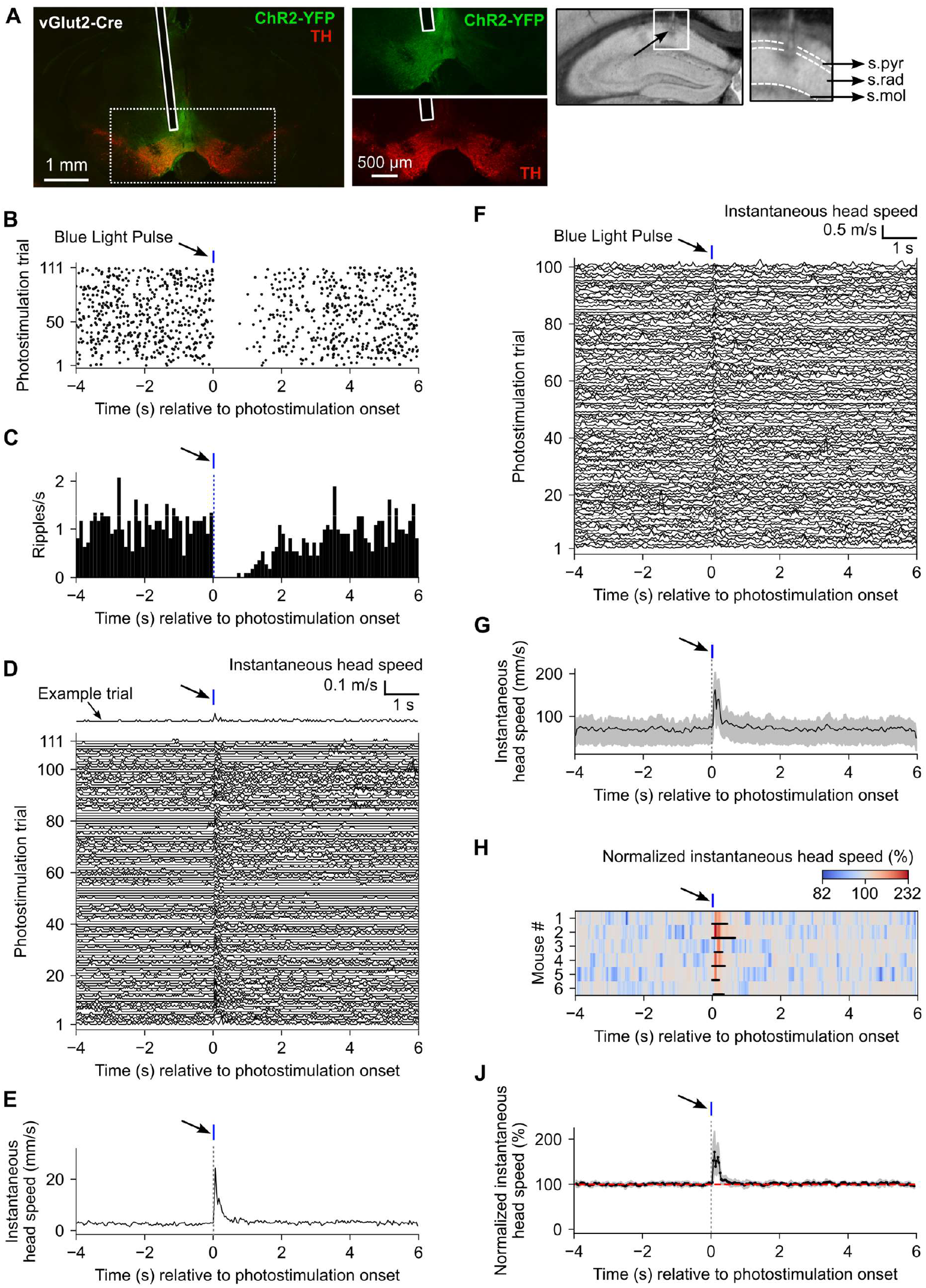
Effect of VTA stimulation on ripple incidence during NREM sleep and on head movement during NREM sleep and wakefulness in vGlut2-Cre mice. **A**) Left: Immunostaining showing overlay of the expression of ChR2-YFP (green) and tyrosine hydroxylase (TH, red) in the VTA of an example mouse. The narrow vertical white rectangle marks the optic fiber track. The green and red channels within the dotted rectangle are expanded in the two adjacent panels. Right: Grayscale images showing the lesion mark of an electrode from the same mouse, with the electrode tip indicated by the black arrow. Data from this electrode are shown in panels B and C. **B**) Raster plot of ripple events (black dots) aligned to onset of a 50-ms stimulation light pulse (blue tick) during NREM sleep. **C**) Peri-stimulus time histogram of ripple rate. **D**) Instantaneous head speed from the same mouse, computed in 33-ms bins for each photostimulation trial and aligned to light-pulse onset during NREM sleep. An example trial is shown on the top. **E**) Trial-averaged head speed from panel D. **F**) Instantaneous head speed from a second example mouse during the awake state, computed in 33-ms bins for each trial and aligned to light-pulse onset. Note the larger scale bar for head speed compared to the NREM sleep (panel D). The last 89 trials are omitted for clarity. **G**) Trial-averaged head speed. Shaded area indicates ± 1 standard deviation (SD). **H**) Pseudocolor plot of normalized instantaneous head speed (%) for the mouse population (n = 6) during the awake state. Red indicates head speeds above baseline; blue indicates head speeds below baseline. Black horizontal bars mark periods of significant modulation in each mouse. **J**) Normalized instantaneous head speed averaged across mice (n = 6) during the awake state. Shaded area indicates ± 1 SD. No significant modulation was detected at the population level.

Beyond the hippocampus, VTA glutamatergic neurons project to multiple targets, raising the possibility that stimulation might produce additional effects. One readily measurable outcome was stimulation-induced change in head position, quantified as instantaneous head speed. The same light pulses that strongly suppressed ripples also caused an abrupt increase in head speed, peaking at ∼20 mm/s and returning to baseline within ∼0.5 s (Fig. 5D, E). During the 1-s period after stimulus onset, the head moved 1.1 ± 0.8 mm (across-trial mean ± 1 standard deviation) away from its mean position (in the 0.1 s) preceding the stimulus. Because vGlut2+ neurons have been implicated in promoting wakefulness, we tested whether this phasic head movement reflected an attempted awakening or a direct motor effect. If the activated vGlut2+ neurons were purely wake-promoting, then stimulation in awake animals should not evoke head movement. Contrary to this prediction, single 50-ms pulses delivered during the awake, ambulatory state elicited robust phasic head movements similar to those observed during NREM sleep (Fig. 5F–J; Fig. S5). Although trial-to-trial variability in the timing of the transient effect (visible as narrow vertical red stripes near time zero in Fig. 5H) precluded a significant population-level change in head speed (n = 6; *p* > 0.05, cluster-size permutation test), the effect was significant in each mouse individually (n = 183–311 trials per mouse; *p* < 0.05, cluster-size permutation test). These results suggest that activation of VTA vGlut2+ neurons can powerfully sup-press hippocampal ripples in NREM sleep and induce phasic head movements in both awake and NREM sleep states, with head movements unlikely to simply reflect a transition to wakefulness. In contrast, activation of GABAergic or dopaminergic neurons induced little (Fig. S4A–D, G, H; n = 6; *p* > 0.05, cluster-size permutation test) or no detectable head movement (Fig. S2A–D, G, H; n = 6; *p* > 0.05, cluster-size permutation test), respectively. Moreover, non-specific activation of VTA neurons in wild-type mice produced prominent head movements (Fig. S6), suggesting that this effect was largely driven by vGlut2+ neurons.

Because our primary goal was to assess the effect of vGlut2+ activation in ripple suppression during sleep, we minimized head movements that, if large enough, could awaken the animal via proprioceptive feedback. For each mouse, we adjusted the blue-light intensity to a level that produced minimal head movement, then conducted the photostimulation experiment with this intensity fixed. In an example mouse, even with moderate light intensity, activation of vGlut2+ neurons (Fig. 6A) produced prominent ripple suppression (Fig. 6B, C) accompanied by minimal head movement (Fig. 6D, E). At the population level (n = 7 mice; six of which are the same as in Fig. 5), single-pulse stimulation significantly suppressed ripples for ∼3 s, with ripple rate returning to near-baseline after ∼6 s (Fig. 6F, G; Fig. S5). Normalized head movement showed a non-significant initial phasic increase followed by a significant ∼3-s sustained increase in instantaneous head speed, also returning to base-line by ∼6 s (Fig. 6H, J). During the 1-s period after stimulus onset, the head moved 0.25 ± 0.2 mm (across-trial mean ± 1 standard deviation) away from its mean position (in the 0.1 s) preceding the stimulus. Green light delivery (intensity matched to the blue light) in the VTA of the same mice, or blue light delivery in a mouse expressing eYFP in VTA vGlut2+ neurons, produced neither ripple suppression nor head movement (Fig. S7). These findings indicate that substantial ripple suppression can be achieved with minimal concurrent head movements.

**Figure 6.**
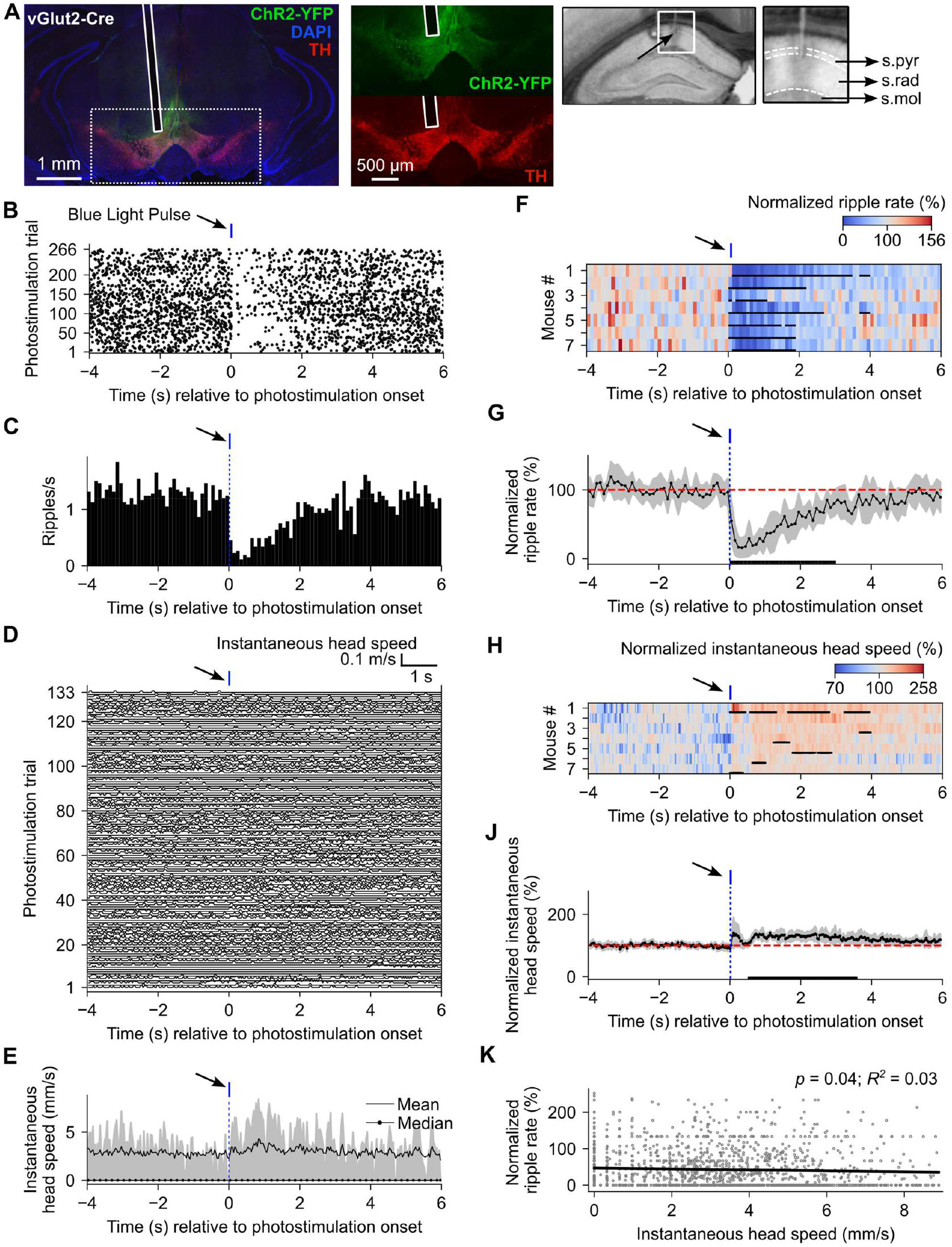
Effect of minimal VTA stimulation on ripple incidence and head movement during NREM sleep in vGlut2-Cre mice. **A**) Left: Immunostaining showing overlay of the expression of ChR2-YFP (green) and tyrosine hydroxylase (TH, red) in the VTA of an example mouse. The narrow vertical white rectangle marks the optic fiber track. The green and red channels within the dotted rectangle are expanded in the two adjacent panels. Right: Gray-scale images showing lesion marks of two electrodes from the same mouse. Data from the electrode with its tip marked by the black arrow are shown in panels B and C. **B**) Raster plot of ripple events (black dots) aligned to onset of a 50-ms stimulation light pulse (blue tick). **C**) Peri-stimulus time histogram of ripple rate. **D**) Instantaneous head speed from the same mouse, computed in 33-ms bins for each photostimulation trial and aligned to light-pulse onset. The last 133 trials are omitted for clarity. **E**) Mean (continuous trace) and median (dotted trace) head speed computed across trials. Shaded area indicates the interquartile range. **F**) Pseudocolor plot of normalized ripple rate (%) for the mouse population (n = 7). Red indicates ripple rates above baseline; blue indicates ripple rates below baseline (suppression). **G**) Normalized ripple rate averaged across mice (n = 7). Shaded area indicates ± 1 standard deviation (SD). **H**) Pseudocolor plot of normalized instantaneous head speed (%) for the mouse population (n = 7). Red indicates speeds above baseline; blue indicates speeds below baseline. **J**) Normalized head speed averaged across mice (n = 7). Shaded area indicates ± 1 SD. **K**) Trial-by-trial correlation between normalized ripple rate and instantaneous head speed (both averaged over the period of significant modulation in panel G), quantified using a linear mixed-effects model fit. The thick dark line indicates the global linear fit (fixed effect). The *p* value and the *R*^*2*^ indicate the significance and variance explained, respectively. In panels F and H, horizontal black bars below each row indicate periods of significant modulation at the individual mouse level. In panels G and J, horizontal black bars indicate significant modulation at the population level.

Although increased head movement accompanied increased ripple suppression on average, it was unclear how reliably these events occurred together. To assess this, we quantified the association between ripple rate and head movement magnitude. For each trial, we calculated the average head speed and the normalized ripple rate within the 3-s window of significant suppression (black bar in Fig. 6G). Ripple rate and head speed showed a significant linear relationship (Fig. 6K; n = 7 mice, 124–389 trials per mouse; *p* = 0.04, linear mixed-effects model), but head speed accounted for only 3% of the variance in ripple rate, indicating a negligible linear association (*R*^*2*^ = 0.03). Given this weak correlation, we next examined ripple suppression in trials with virtually no head movement following VTA stimulation (Fig. 7A, B). Even in these trials, ripple suppression remained significant (Fig. 7C, D), lasting ∼2 s after stimulation. These findings indicate that ripple suppression occurs largely independently of detectable head movement.

**Figure 7.**
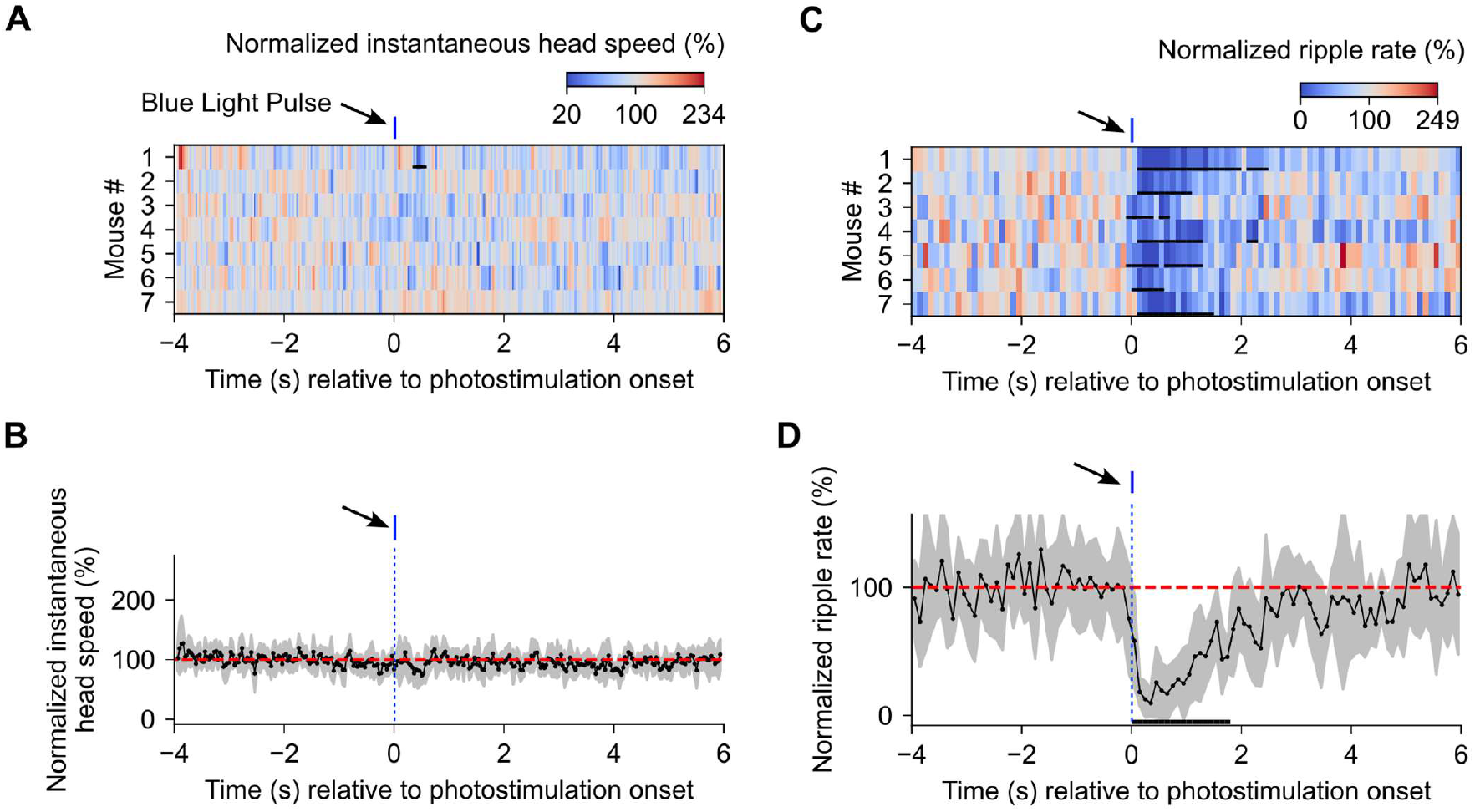
Effect of minimal VTA stimulation on ripple incidence during NREM sleep in vGlut2-Cre mice in trials with negligible stimulus-induced head movements. **A**) Pseudocolor plot of normalized instantaneous head speed (%) for the mouse population (n = 7). Red indicates speeds above baseline; blue indicates speeds below baseline. The blue tick above the plot marks the 50-ms blue-light pulse. Black horizontal bar marks a period of significant modulation in mouse 1. **B**) Normalized head speed averaged across mice (n = 7). Shaded area indicates ± 1 standard deviation (SD). No significant modulation was detected at the population level. **C**) Pseudocolor plot of normalized ripple rate (%) for the mouse population (n = 7). Red indicates ripple rates above baseline; blue indicates ripple rates below baseline (suppression). Black horizontal bars mark periods of significant modulation in each mouse. **D**) Normalized ripple rate averaged across mice (n = 7). Shaded area indicates ± 1 SD. The black horizontal bar marks the period of significant modulation across the population.

## Discussion

While sharp-wave ripples have been characterized extensively for their potential role in memory consolidation, the mechanisms regulating their occurrence are only beginning to be understood. Using cell-type specific and temporally precise optogenetic stimulation, we found that activation of glutamatergic neurons in the VTA strongly suppressed hippocampal sharp-wave ripples. In addition, largely independent of the ripple suppression, activation of these neurons also produced prominent head movements during both sleep and awake states. These results identify a potential approach to reduce the ripple incidence and motivate future studies to dissect the role of VTA glutamatergic neurons in movement.

In exploring the influence of subcortical regions on hippocampal ripples, previous studies have demonstrated ripple suppression using various neuronal activation protocols. For example, Novitskaya et al. [17] used brief (∼100–200 ms), high-frequency (∼100 Hz) electrical stimulation of the locus coeruleus; Vandecasteele et al. [18] employed prolonged (1–60 s) sinusoidal (1–12 Hz) photostimulation of the medial septal cholinergic neurons; and Wang et al. [16] applied brief (2 × 3 ms) photostimulation of median raphe neurons in wild-type mice. Although the prolonged stimulation used by Vandecasteele et al. precludes direct comparison with our results, the effectiveness of brief stimulation in suppressing ripples in the locus coeruleus [17] and median raphe [16] is comparable to our findings in the VTA, where brief photostimulation of VTA neurons in wild-type mice or selectively in glutamatergic neurons produced strong ripple suppression that outlasted the stimulus.

An unexpected outcome of VTA stimulation was head movement, which occurred in parallel with ripple suppression. Previous studies reported ripple suppression through subcortical neuronal activation during sleep [16, 17], but they did not describe movement time-locked to the stimulation. By contrast, studies focused on VTA’s role in movement—but not ripple suppression—have shown that DA and GABAergic neurons encode specific movement parameters during the awake state. Using a head-fixed system that measured forces during behavior, Hughes et al. [32] showed that VTA dopaminergic neurons represent the impulse vector (force generated over time) of forward and backward movements, and that photoactivation of DA neurons with either a single 5-ms light pulse or a pulse train (3–12 pulses at 25 Hz) elicited net forward movement. In our experiments, however, photoactivation of the DA neurons with either a single 50-ms pulse or a pulse train (50 × 50 ms) during NREM sleep produced no detectable head movement, suggesting that DA neuron function may be strongly state dependent. Similarly, VTA GABAergic neurons have been shown in awake mice to encode principal head angles (pitch, yaw, and roll) and forward/backward force vectors; their optogenetic activation alters head orientation [34] and suppresses forward force exertion [33]. Although our experimental design does not permit a direct comparison with these awake-state findings, during NREM sleep we observed only small head movements following GABAergic stimulation, again suggesting behavioral-state dependence. In contrast, the role of VTA glutamatergic neurons in movement has not been described previously. Here we show that their activation produces robust head movements in both NREM sleep and awake states, motivating further investigation.

A previous study reported that optogenetic activation of VTA vGlut2+ neurons during NREM sleep promoted wakefulness [35] using a prolonged stimulation protocol (60 s at 20-Hz, totaling 1200 pulses). In contrast, the brief (50 ms), single-pulse photostimulations used in our study were not sufficient to induce a transition from NREM sleep to wakefulness. Moreover, because photostimulation evoked head movements both during NREM sleep and when the mice were actively moving in their home cages, the small head movements we observed are unlikely to reflect sleep-wake transitions. Instead, these findings suggest that VTA vGlut2+ neurons may play a distinct role in regulating head movement.

Vagus nerve stimulation (VNS) is an FDA-approved treatment for several disorders, including epilepsy, stroke, and depression [36]. A recent study demonstrated that VTA neurons are activated in rats trained to self-administer VNS via lever pressing [37], suggesting that VNS could provide a minimally invasive means to recruit VTA activity. Because sharp-wave ripples are critical for memory consolidation, an intriguing possibility is that VNS-mediated activation of VTA during NREM sleep could suppress ripples and thereby weaken pathological memories, such as those associated with PTSD. Future studies will be needed to test this potential therapeutic approach.

Our study has several limitations. First, we did not selectively express ChR2 in VTA neurons projecting exclusively to the hippocampus, and thus our stimulation likely recruited multiple VTA projection targets in addition to the hippocampus. Future studies will be needed to dissect the relative contribution of these different pathways to the ripple suppression effect. Nevertheless, our finding that VTA activation can suppress ripples suggests a potential avenue for altering memory consolidation for therapeutic purposes, regardless of whether the effect is mediated through direct or indirect projections. Second, our movement analysis was restricted to the head, as tracking was based on an LED attached to the microdrive implant. We therefore cannot determine whether activation of the glutamatergic neurons also elicited movements of other body parts. Moreover, because our overhead camera tracked a single LED, we could not resolve pitch, yaw, and roll components of head movement. Future studies using higher-resolution approaches, such as those employed by Hughes et al. [32] will be necessary to characterize the detailed kinematics of VTA glutamatergic neuron-evoked movements.

## Conclusion

The VTA is a neuronal hub composed of multiple cell types with diverse afferent and efferent projections [29, 38, 39]. To minimize the pleiotropic effects expected from broad VTA activations, we employed brief, temporally precise, cell type-specific stimulations. We found that activation of VTA glutamatergic neurons suppressed hippocampal sharp-wave ripples during NREM sleep. Notably, non-specific activation of the VTA neurons produced a comparable level of ripple suppression. Thus, under therapeutic contexts where cell-type-specific targeting remains a major challenge, our results suggest that non-specific VTA activation may be used therapeutically to suppress traumatic memories. Beyond its well-known role in reward processing, the VTA has been implicated in a wide range of functions. Our observation that glutamatergic neuronal activation elicits brief head movements during both wakefulness and NREM sleep adds to this growing body of work and highlights a potential role for VTA glutamatergic neurons in movement control requiring further investigation.

## Methods

### Animals

Wild type C57BL/6J and knock-in mice (DAT-Cre, vGAT-Cre, and vGlut2-Cre) were purchased from the Jackson Laboratory (DAT-Cre, Strain #:006660; vGAT-Cre, Strain #: 028862; vGlut2-Cre, Strain #:028863). They were housed under a 12-h light/dark cycle with food and water available ad libitum. Mice were group-housed until they were at least 2 months old, after which they were single-housed following implantation of micro-drives. Both male and female mice were used for experiments. All procedures and animal care were carried out in compliance with the guidelines specified by the Institutional Animal Care and Use Committee at the University of Pennsylvania.

### Microdrive assembly

We custom-designed and 3D-printed microdrives carrying a grid of 4–6 depth-adjustable tetrodes for recording. Tetrodes were made by twisting four insulated Nichrome wires (California Fine wires, Stablohm 800A, material #100189, 12.7 µm core, 20 µm outer diameter, heavy polyimide coated). The four tetrode terminals were shorted together and gold plated to an impedance of 100 – 300 KΩ, creating a single recording channel. Ground and reference silver wires (A-M Systems), along with the tetrodes, were connected to a light-weight electrode interface board (modified EIB-18, Neuralynx Inc.). The tetrodes were carried on a shuttle movable by screws. The assembled microdrive weighed < 1g. For a subset of mice, we used a non-adjustable version of this microdrive.

### AAV virus stocks

For optogenetics experiments, we used concentrated viral suspensions of AAV5-EF1a-DIO-hChR2(H134R)-eYFP and AAV5-EF1a-DIO-eYFP (4 × 10^12^ viral particles/mL), prepared by the University of North Carolina Vector Core Facility.

### Surgical procedures

In each mouse, two surgeries were performed: the first for vital infusion into the VTA and the second, 3–4 weeks later, for implanting both the microdrive in the hippocampus and an optic fiber in the VTA (Fig. 1A). After anesthesia induction with 3% isoflurane, mice were transferred to a stereotaxic frame, where isoflurane was maintained at 1.75– 2.25% (flow rate 1 L/min). For viral infusion, a burr hole was drilled at AP: −3.16 mm and ML: ± 0.4 mm (Franklin & Paxinos, 2008), and 0.3 µL of viral suspension was delivered at 0.1 µL/min to DV: −4.24 mm. For microdrive implantation, a burr hole was drilled at AP: −1.94 mm, ML: ± 1.18 mm for inserting the electrode array. The ground wire was attached to a screw (Antrin Miniature Specialists) over the right cerebellum; the reference wire’s silver ball tip was placed on the brain surface via a burr hole over the left frontal bone. For depth-adjustable microdrives, electrodes were lowered to DV: −1.00 mm during surgery and advanced to the final target 2–3 days later. For non-adjustable microdrives, the electrodes were placed at the final depth of DV: −1.25 mm during surgery. In addition, a chronic optic fiber (numerical aperture, 0.48; core diameter, 200 µm; Thorlabs) was inserted through a burr hole at a 6^°^ angle, targeting the fiber tip at AP: −3.16mm, ML: ± 0.35 mm, DV: −3.85 mm. The microdrive and the optic fiber were secured to the skull using C & B MetaBond (Parkell) dental cement.

### Experimental setup

We recorded AC-coupled broadband (0.1 Hz–16 KHz) neural signals continuously at 32 KHz sampling rate using a Neuralynx data acquisition system (Digital Lynx SX) with a unity-gain headstage preamplifier (HS-18). A red LED on the preamplifier enabled mouse position tracking. For optical stimulation, a 450 nm or 561 nm laser (Changchun New Industries) was connected to VTA-implanted optic fiber via a lightweight patch cord (Doric Lenses) and paired with a lightweight electrical recording tether (Conner wire). Commutators on both cables prevented tangling. TTL pulses from a Master-9 stimulator (AMPI) controlled laser light timing and duration and were time-stamped in the Neuralynx system for alignment with neural recordings.

### Recording procedure

After microdrive implantation, animals recovered for at least two days before electrodes were lowered ∼80 µm/day until sharp-wave ripples were detected during NREM sleep. Mice with non-adjustable electrodes were left undisturbed for 3–4 days to recover and allow the electrodes to stabilize. Before experiments, mice were habituated for ≥3 days to the recording setup, sleeping in their home cage with both the electrical tether and optic fiber patch cord attached (Fig. 1B). Experiments were conducted during the light phase, a median of 4 h (IQR: 3–6 h) after light onset. During recordings, mice remained in their home cage inside a sound-attenuating enclosure (Med Associates). To prevent cable tangling, the water bottle from the mouse cage was removed and food pellets were moistened to reduce dehydration. To keep mice from climbing out, another cage with its bottom removed was stacked on top of the home cage. Mouse movement was monitored continuously with an overhead video camera (30 frames/s) synchronized with neural recordings.

### Optical stimulation experiments

With continuous recording of dorsal hippocampal neural activity and video tracking of mouse position, we optically activated the VTA in multiple stimulation “trials” (Fig. 1C). Each stimulation trial consisted of VTA stimulation—either a single light pulse or a pulse train—followed by a variable period without stimulation. In wild-type mice, the VTA was stimulated with either a single 5-ms pulse (n = 2 mice) or a pair of 3-ms pulses separated by 40 ms inter-pulse interval (n = 1). In all other mice, 50-ms light pulses were used. For pulse-train trials, 50 pulses were delivered with an inter-pulse interval of 100 ms. The inter-trial intervals were 15–20 s for single-pulse trials and 35–40 s for pulse-train trials. The intensity of both blue and green lights at the tip of the implanted optic fiber was 9.8 ± 1.5 mW/mm^2^. For the experiments in Figure 6, where light intensity was titrated for each mouse, the median intensity was 1.1 mW/mm^2^ (IQR: 0.9–7.8). Stimulation was applied only during the intended behavioral state (sleep or awake), which was determined by animal movement and field-potential oscillations characteristic of each state.

### Head movement analyses

To compute the instantaneous head speed, we detected the mouse’s position in each video frame online by thresholding the LED tracker light. Offline, we calculated the frame-to-frame position displacement in pixels, converted it to millimeters (mm), and divided by the video-frame duration (0.033 s) to obtain instantaneous speed in mm/s. To quantify the largest distance the head moved in response to photostimulation, in each trial, we calculated the maximum distance of the head from its mean position during the 0.1 s preceding the stimulus, within the 1-s post-stimulus period. To correct for slow head drifts, we computed the maximum displacement during a 1-s baseline period and subtracted this value from the post-stimulus head displacement. Corrected displacements were then averaged across trials and subsequently across mice.

### Identification of behavioral states

We identified behavioral states based on mouse motion and the theta/delta ratio (TDR)—ratio of the spectral power in the hippocampal theta (4–11 Hz) and delta (1–4 Hz) bands, as previously described [40]. Briefly, time periods with motion values below a threshold *δ*_*m*_ were classified as putative sleep episodes. Within these, segments with TDR moving-variance below a threshold *δ*_*NREM*_ were labeled as putative NREM states, and those with TDR moving-variance above a threshold *δ*_*REM*_ were labeled as putative REM states. To refine these classifications, we removed spurious segments defined as those shorter than 30s or separated by gaps exceeding 3 min between adjacent NREM and REM segments, yielding confirmed sleep states. The thresholds *δ*_*m*_, *δ*_*REM*_, and *δ*_*REM*_ were determined heuristically and applied uniformly across all mice. Periods outside putative sleep episodes were treated as awake states.

### Detection of ripples

Ripple events were detected in each electrode as threshold-crossing events, as previously described [6], with modifications. Specifically, we band-pass filtered the raw data (Fig. 1D, top panel) between 120 and 250 Hz (transition band: 25 Hz; Fig. 1D, middle panel) and detected ripples in 2-s contiguous segments using the following steps: *1*) Squared the filtered signal (gray trace in Fig. 1D, bottom panel) and computed its standard deviation σ; *2*) Computed the filtered signal’s envelope by detecting peaks, interpolating between adjacent peaks, and smoothing the result with a Gaussian window (dark continuous trace in Fig. 1D, bottom panel); *3*) Detected peaks in the envelope and selected as ripple events those that crossed a threshold of 9σ, had a width of at least 30 ms (measured at 50% peak prominence), and were separated by >50 ms from adjacent peaks. The timing of the selected peaks corresponded approximately to the maximum amplitude of the ripple oscillations.

### Analysis of the effects of VTA stimulation

To examine the effects of VTA stimulation on ripple occurrence and instantaneous head speed, trials were aligned to stimulation onset, and a pre-stimulus baseline and post-stimulus window were defined. For single- or two-pulse stimulation, a 4-s baseline and a 6-s post-stimulus period were used. For 50-pulse trains, a 10-s baseline and 15-s post-stimulus period were used to allow effects of the longer stimulation to dissipate before the next trial. Ripple events were binned in 100 ms intervals to calculate ripple rate (ripples/s) for each bin, then averaged across trials. In mice with multiple electrodes, ripple rates in a given bin were first averaged across electrodes within each trial before averaging across trials. Instantaneous head speed was binned in 33.3 ms intervals and averaged across trials. To average these measures across mice, trial-averaged values were normalized by dividing by the mean baseline value and multiplying by 100.

To examine the effect of VTA stimulation on ripple rate without confounding head movements, we selected photostimulation trials in which post-stimulus head movement was comparable to baseline. For each trial, we computed the ratio of average post-stimulus head speed to pre-stimulus head speed and selected trials with ratios between 0.5–1.0. Because the stimulation-evoked head movements were transient and could be diluted in the overall post-stimulus average, we weighted the initial 275 ms of the post-stimulus period—when most transient movement occurred—approximately seven times higher than an equivalent duration of the remaining post-stimulus period during speed averaging.

### Histology

To localize the electrode tips, we anesthetized each mouse with a ketamine-xylazine mixture and passed 30 µA positive current for 8–10 s through the electrode tips, using the ground or reference electrode as the return path. We then performed cardiac perfusion using 10% formalin (in phosphate-buffered saline), left the mouse head with the microdrive implant in the fixative at 4 °C for one or more days, removed the brain, placed it in the fixative an additional day, and sliced it coronally at 50 µm thickness using a vibratome.

### Immunofluorescence

We stained brain slices for tyrosine hydroxylase (TH) and GFP using the following procedure: *1*) Incubated the slices in a blocking solution (10% horse serum, 0.2% BSA, 0.5% Triton-X) for 2 h at room temperature; *2*) Transferred the slices to a carrier solution (1% horse serum, 0.2% BSA, 0.5% Triton-X) containing primary antibodies against TH (rabbit anti-TH, 1:1000 dilution, Millipore) and GFP (mouse anti-GFP, 1:1000 dilution, Thermo Fisher Scientific) and incubated overnight at 4 °C; *3*) Washed the slices four times with phosphate buffered saline (PBS); *4*) Transferred the slices to the carrier solution containing secondary antibodies (goat anti-Rabbit-Alexa 594 and anti-mouse-Alexa 488, 1:1000 dilution, Thermo Fisher Scientific), incubated for 2 h, and washed four times with PBS. The stained slices were mounted using anti-fade mounting medium and images were acquired using Olympus BX63 automated fluorescence microscope.

### Exclusions

Eight mice from our dataset were excluded due to lack of detectable ripples in the recordings (n = 4), misplacement of the optic fiber outside the VTA (n = 1), or technical errors (n = 3).

### Statistical analysis

At both single-mouse and population levels, we assessed the statistical significance of light stimulation-induced changes in ripple rate and head movement speed using nonparametric tests as previously described [41]. To apply these tests, we restricted the post-stimulus period to match the baseline duration. The goal was to identify contiguous post-stimulus time bins where ripple rate or head speed differed significantly from baseline while accounting for multiple comparisons across time bins. At the single-mouse level for ripple rate, the procedure was the following: *1*) For each of “*n*” trials, mean ripple rate across the baseline period was computed and paired-sample t-test was performed comparing these *n* baseline averages to the corresponding *n* ripple rates of each of the “*m*” post-stimulus bins, yielding *m* t-values; *2*) Contiguous bins with significant modulation (*p* < 0.05) were grouped into clusters, and each cluster’s “cluster mass” was calculated by summing its t-values; *3*) A null distribution of cluster masses was created by randomly swapping baseline and post-stimulus periods within each trial with 0.5 probability, repeating steps 1 and 2, 2000 times, and recording the largest cluster mass in each iteration; *4*) The *p*-value for each observed cluster was computed as the fraction of null distribution values exceeding that cluster’s mass. At the mouse-population level, the same procedure was applied using trial-averaged ripple rate from each mouse instead of individual trial data. The same method was used to assess significance of stimulation-induced modulation of head speed. Clusters with *p* < 0.05 were considered significantly modulated.

To compute the correlation between ripple rate and head speed, we first identified the post-stimulus time period showing significant ripple suppression at the mouse-population level. For each trial, we computed the mean ripple rate during this period and normalized it by dividing by the trial’s baseline mean ripple rate and multiplying the result by 100. Similarly, we computed the mean instantaneous head speed during the same post-stimulus period for each trial. We then fitted a linear mixed-effects model (using the *Statsmodels* Python package), with mean instantaneous head speed as the fixed effect, normalized ripple rate as the response variable, and mouse identity as a random effect. The variance explained by the model was quantified using the conditional *R*^2^ metric [42], which accounts for mouse-to-mouse variation in ripple rate and head-speed ranges.

## Acknowledgements

We thank Marion Scott for technical assistance. This work was supported by National Institute on Drug Abuse Grant R37 DA053296 and by a generous award from the Chernowitz Medical Research Foundation.

## Author contributions

M.S conceived the study, performed surgeries, conducted experiments, analyzed data, and wrote initial draft of the manuscript. S.M prepared micro-drives, performed surgeries, conducted experiments, and carried out histology, immunohistochemistry, and imaging. M.S.Z performed surgeries, histology, immunohistochemistry, and imaging. J.A.D conceived the study, provided funding, resources, and supervision, and edited the manuscript. All authors reviewed and approved the final manuscript.

## Data availability

All data supporting the findings of this study are included in the manuscript and its supplementary materials. Raw data are available from the corresponding author upon reasonable request.

## Supplementary Information

**Figure S1.**
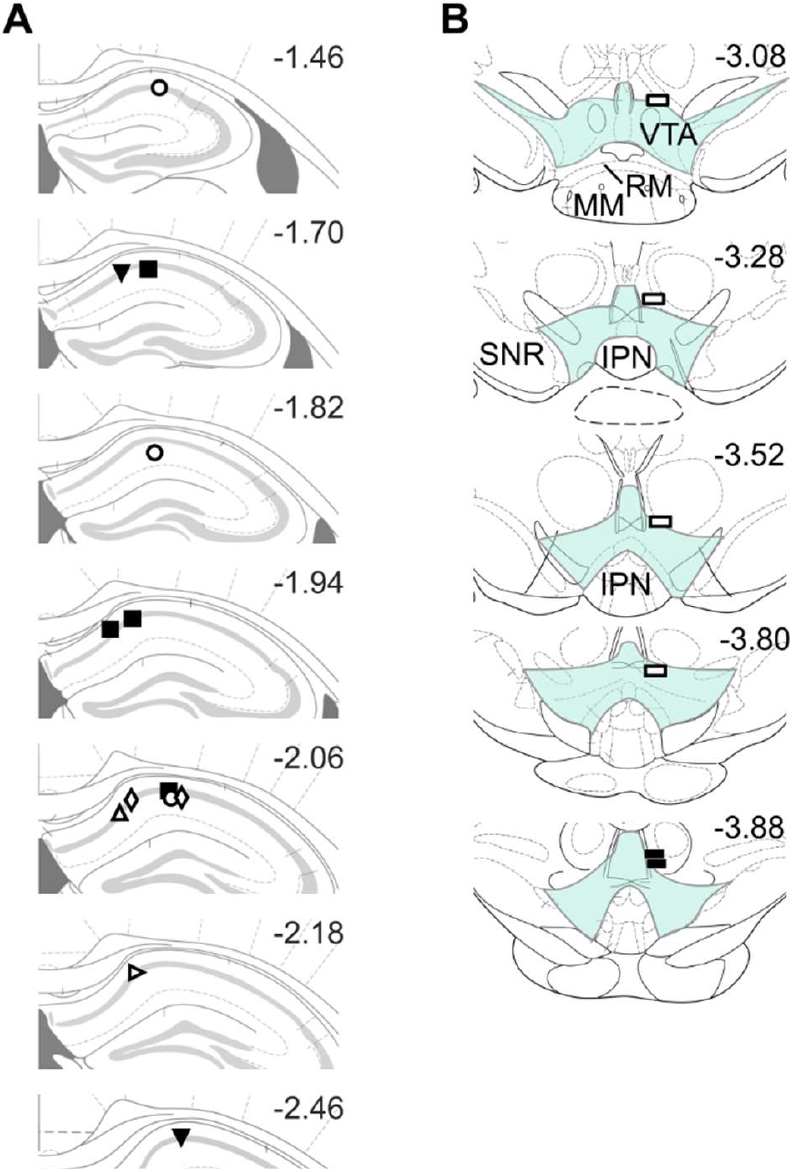
Electrode and optic fiber locations in DAT-Cre mice. **A**) Hippocampal electrode tip locations, with marker type distinguishing mice. **B**) Optic fiber tip locations (rectangles). Blue shaded region indicates the VTA, with nearby structures labeled (IPN, interpeduncular nucleus; MM, medial mammillary nuclei; RM, retromammillary nucleus; SNR, substantia nigra pars reticulata). In both panels (A, B), fill colors indicate sex (black = male, white = female).

**Figure S2.**
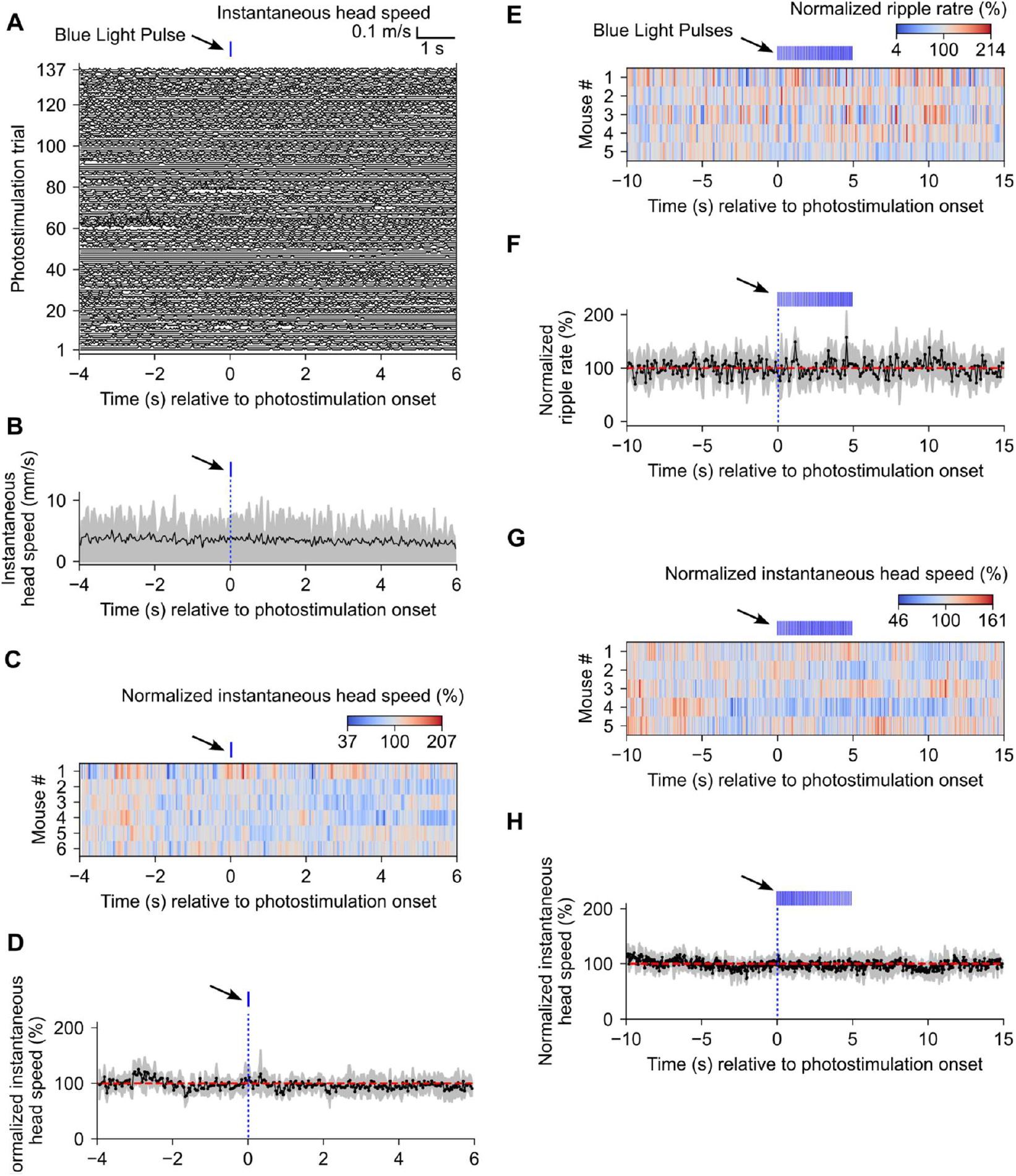
Effect of VTA stimulation on head movement and ripple incidence during NREM sleep in DAT-Cre mice. **A**) Instantaneous head speed computed in 33-ms bins for each photostimulation trial and aligned to light-pulse onset (blue tick above) for the same example mouse shown in Fig. 3A-C. **B**) Trial-averaged head speed from panel A. Shaded area indicates the interquartile range. **C**) Pseudocolor plot of normalized instantaneous head speed (%) across the mouse population (n = 6; same as in Fig. 3D). Red indicates speeds above baseline; blue indicates speeds below baseline. **D**) Normalized head speed averaged across mice (n = 6). Shaded area indicates ± 1 standard deviation (SD). **E**) Pseudocolor plot of normalized ripple rate (%) for a subset of the mice (n = 5) from the population in panel C. The blue ticks above the plot mark the 50 × 50-ms blue-light pulse train. Red indicates ripple rates above baseline; blue indicates ripple rates below baseline (suppression). **F**) Normalized ripple rate averaged across mice (n = 5). Shaded area indicates ± 1 SD. **G**) Pseudocolor plot of normalized instantaneous head speed (%) for the mouse population (n = 5). Red indicates speeds above baseline; blue indicates speeds below baseline. **H**) Normalized head speed averaged across mice (n = 5). Shaded area indicates ± 1 SD. No significant modulation was detected either at the level of individual mice (panels C, E, and G) or at the population level (panels D, F, and H).

**Figure S3.**
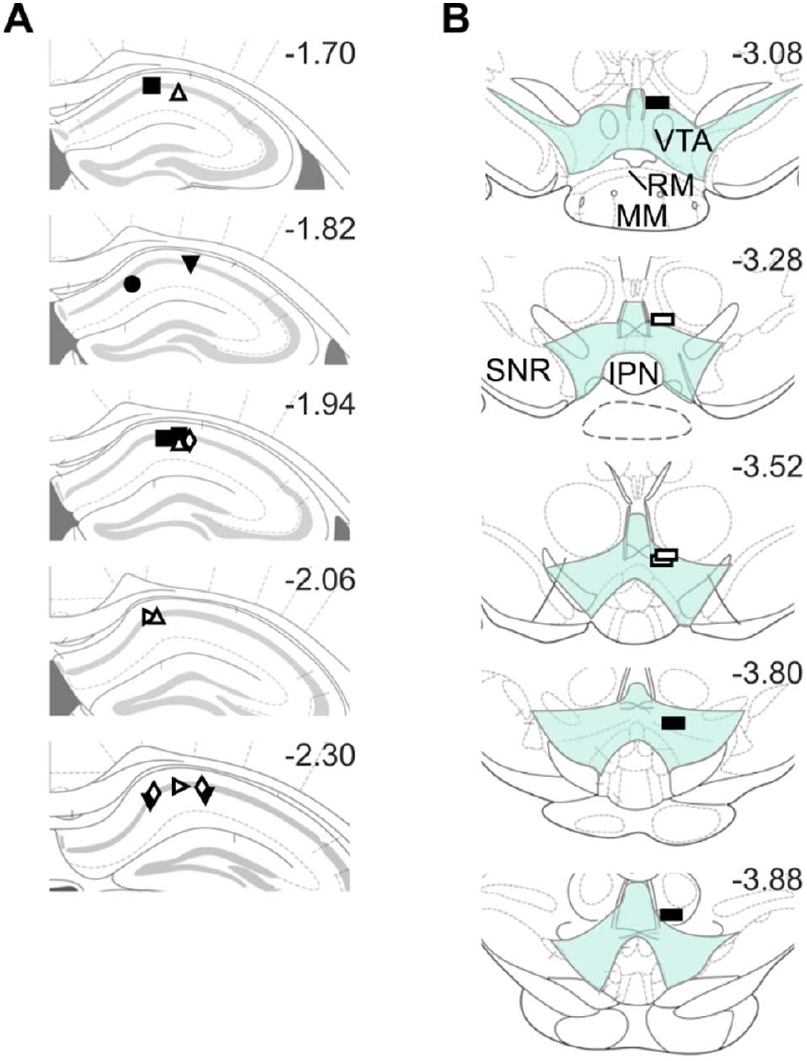
Electrode and optic fiber locations in vGAT-Cre mice. A) Hippocampal electrode tip locations, with marker type distinguishing mice. B) Optic fiber tip locations (rectangles). Blue shaded region indicates the VTA, with nearby structures labeled (IPN, interpeduncular nucleus; MM, medial mammillary nuclei; RM, retromammillary nucleus; SNR, substantia nigra pars reticulata). In both panels (A, B), fill colors indicate sex (black = male, white = female).

**Figure S4.**
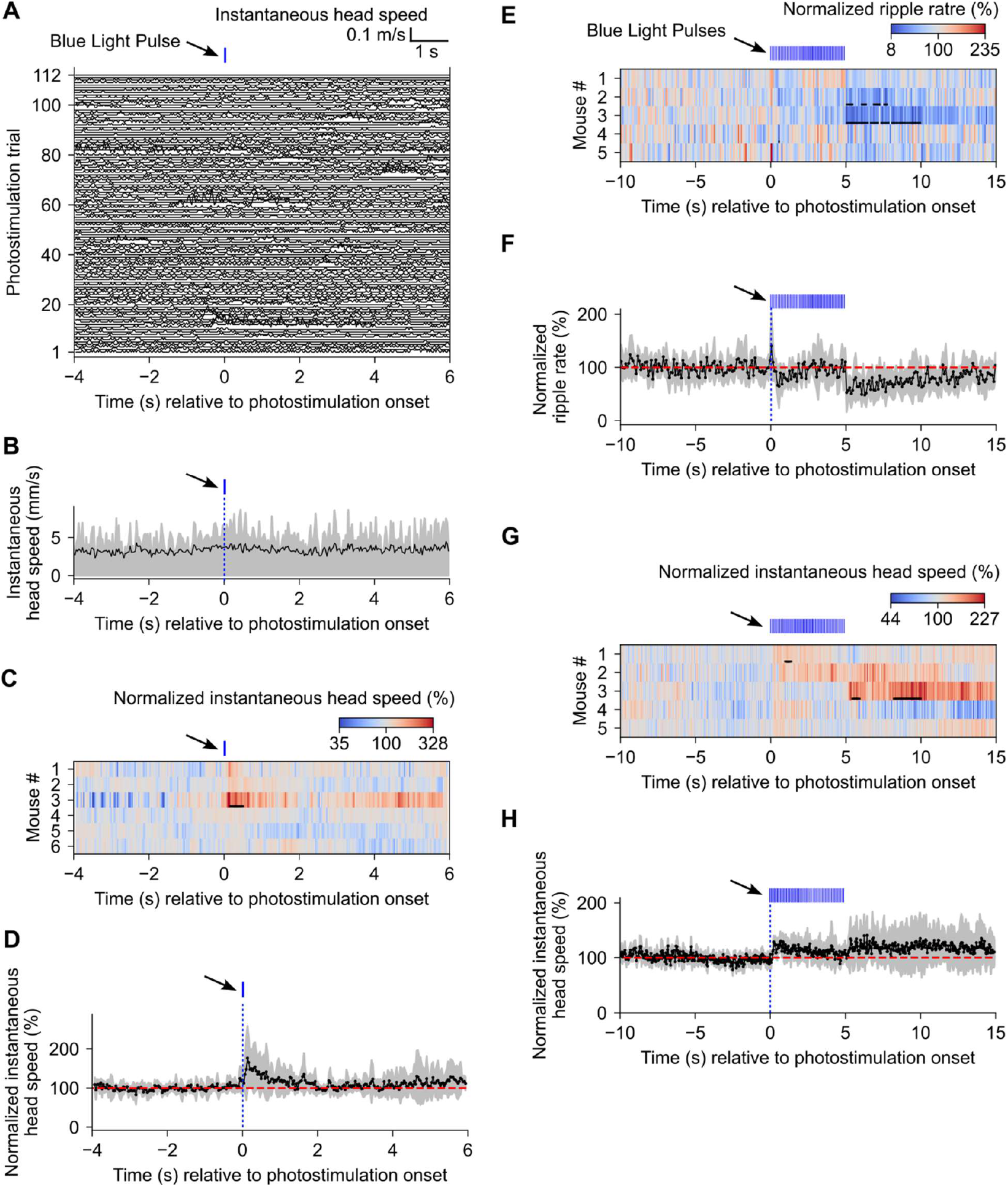
Effect of VTA stimulation on head movement and ripple incidence during NREM sleep in vGAT-Cre mice. **A**) Instantaneous head speed computed in 33-ms bins for each photostimulation trial and aligned to light-pulse onset (blue tick above) for the same example mouse shown in Fig. 4A-C. The last 113 trials are omitted for clarity. **B**) Trial-averaged head speed. Shaded area indicates the interquartile range. **C**) Pseudocolor plot of normalized instantaneous head speed (%) across the mouse population (n = 6; same as in Fig. 4D). Red indicates speeds above baseline; blue indicates speeds below baseline. **D**) Normalized head speed averaged across mice (n = 6). Shaded area indicates ± 1 standard deviation (SD). **E**) Pseudocolor plot of normalized ripple rate (%) for a subset of the mice (n = 5) from the population in panel C. The blue ticks above the plot mark the 50 × 50-ms blue-light pulse train. Red indicates ripple rates above baseline; blue indicates ripple rates below baseline (suppression). **F**) Normalized ripple rate averaged across mice (n = 5). Shaded area indicates ± 1 SD. **G**) Pseudocolor plot of normalized instantaneous head speed (%) for the mouse population (n = 5). Red indicates speeds above baseline; blue indicates speeds below baseline. **H**) Normalized head speed averaged across mice (n = 5). Shaded area indicates ± 1 SD. In panels C, E, and G, horizontal black bars below each row indicate periods of significant modulation at the individual mouse level. In panels D, F, and H, no significant modulation at the population level was detected.

**Figure S5.**
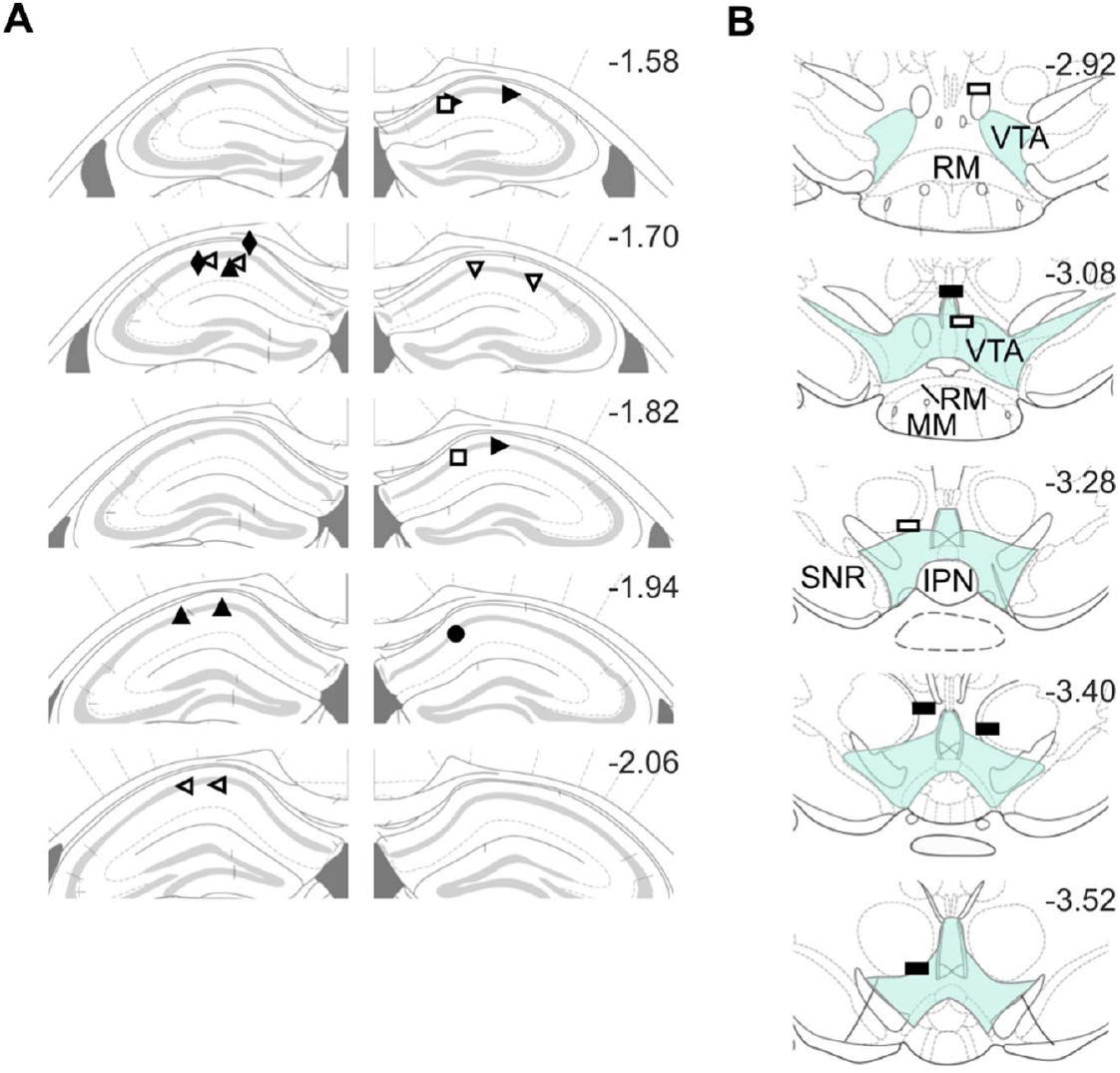
Electrode and optic fiber locations in vGlut2-Cre mice. **A**) Hippocampal electrode tip locations, with marker type distinguishing mice. **B**) Optic fiber tip locations (rectangles). Blue shaded region indicates the VTA, with nearby structures labeled (IPN, interpeduncular nucleus; MM, medial mammillary nuclei; RM, retromammillary nucleus; SNR, substantia nigra pars reticulata). In both panels (A, B), fill colors indicate sex (black = male, white = female).

**Figure S6.**
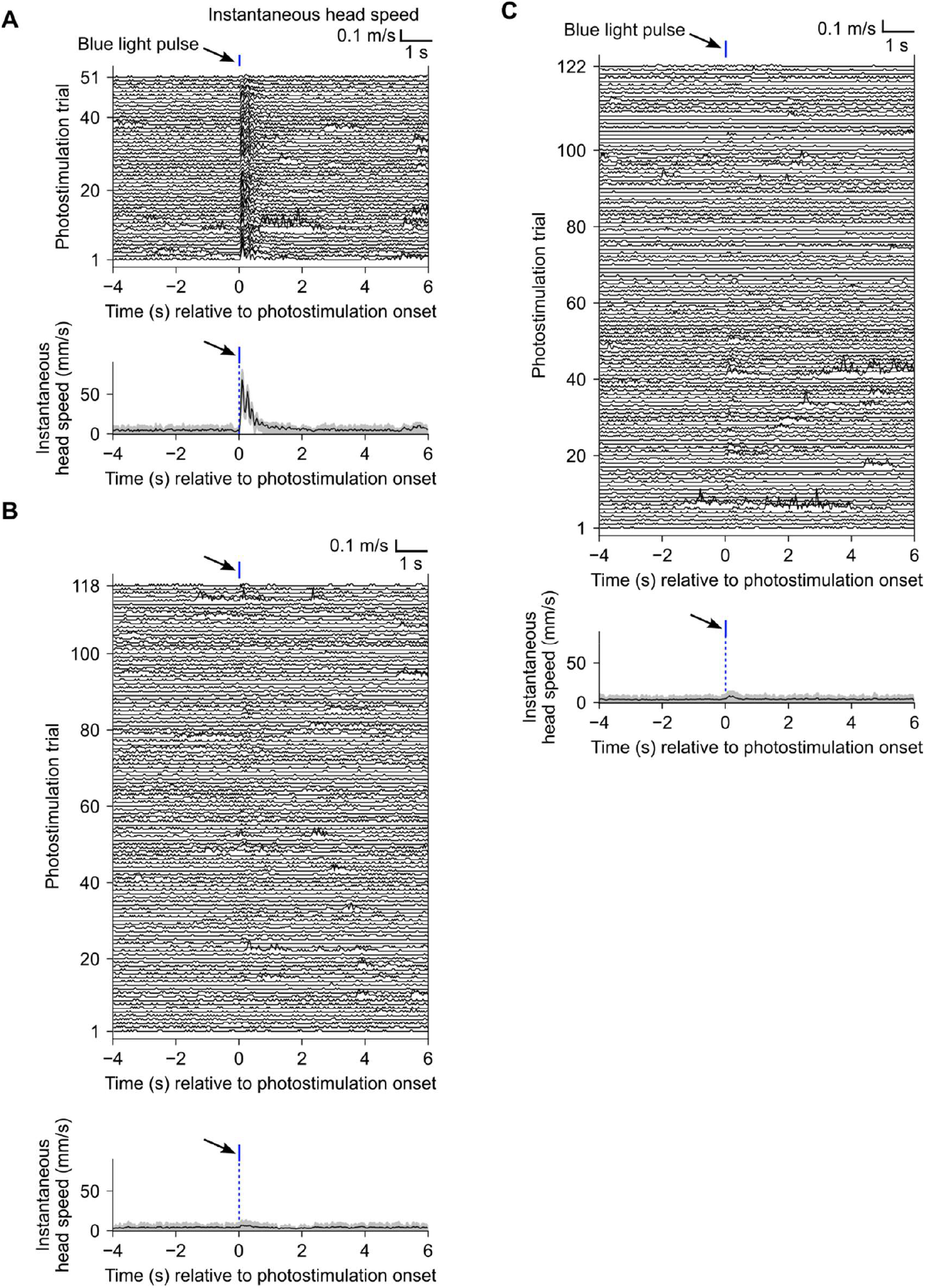
Instantaneous head speed in individual wild-type mice. Panels A–C correspond to three different mice. **A**–**C**) Top: Instantaneous head speed computed in 33-ms bins for each photostimulation trial and aligned to light-pulse onset (blue tick above). Bottom: Trial-averaged head speed from the corresponding top panel. In panel C, the last 122 trials are omitted for clarity. Shaded regions indicate the interquartile range.

**Figure S7.**
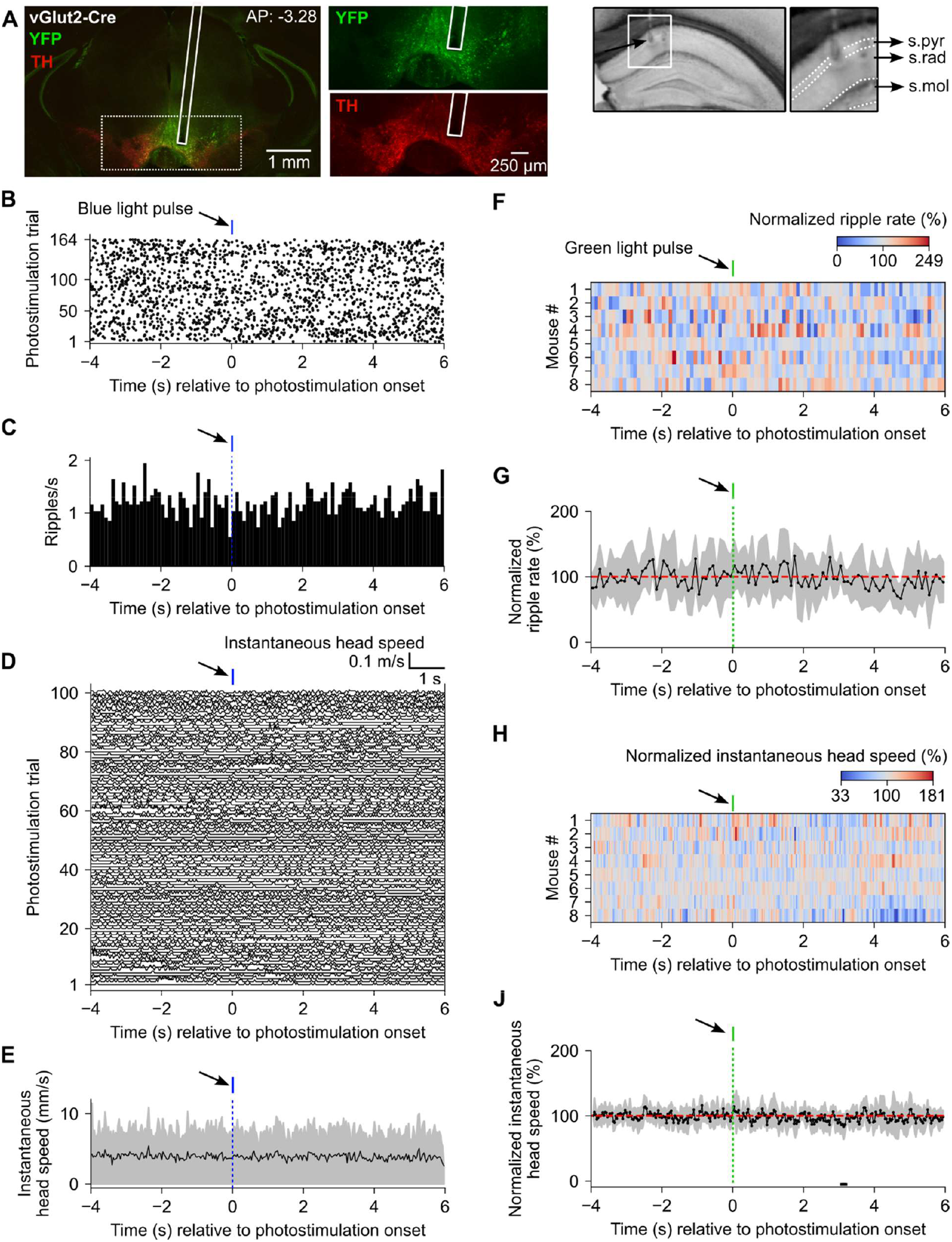
Control experiments for effects observed in vGlut2-Cre mice. **A**) Left: Immunostaining showing overlay of the expression of YFP (green) and tyrosine hydroxylase (TH, red) in the VTA of an example vGlut2-Cre mouse. The narrow vertical white rectangle marks the optic fiber track. The green and red channels within the dotted rectangle are expanded in the two adjacent panels. Right: Grayscale images showing lesion marks of two electrodes from the same mouse. Data from the electrode with tip marked by the black arrow are shown in panels B and C. **B**) Raster plot of ripple events (black dots) aligned to onset of a 50-ms blue light pulse (blue tick). **C**) Peri-stimulus time histogram of ripple rate. **D**) Instantaneous head speed from the same mouse, computed in 33-ms bins for each photostimulation trial and aligned to light-pulse onset. The last 64 trials are omitted for clarity. **E**) Trial-averaged head speed. Shaded area indicates the interquartile range. **F**) Pseudocolor plot of normalized ripple rate (%) for the mouse population [n = 8 (6 vGlut2-Cre mice, same as in Fig. 6; 2 wild-type mice, same as in Fig. 2)]. Red indicates ripple rates above baseline; blue indicates ripple rates below baseline (suppression). The green tick above the plot marks the 50-ms green light pulse. **G**) Normalized ripple rate averaged across mice (n = 8). Shaded area indicates ± 1 standard deviation (SD). **H**) Pseudocolor plot of normalized instantaneous head speed (%) for the mouse population (n = 8). Red indicates speeds above baseline; blue indicates speeds below baseline. **J**) Normalized head speed averaged across mice (n = 8). Shaded area indicates ± 1 SD. No significant modulation was detected either at the individual mouse level (panels F and H) or at the population level (panels G and J).

